# Tris inhibits a GH1 β-glucosidase by a linear mixed inhibition mechanism

**DOI:** 10.1101/2024.09.28.615569

**Authors:** Rafael S. Chagas, Sandro R. Marana

**Author notes:** Corresponding author. Sandro R. Marana.

## Abstract

Here we demonstrate that Tris (2-amino-2-(hydroxymethyl)-1,3-propanediol), largely used as buffering agent, is a linear mixed inhibitor (*K*_i_ = 12 ± 2 mM and α = 3 ± 1) of the GH1 β-glucosidase from the insect *Spodoptera frugiperda* (Sfβgly). Such inhibition mechanism implies in the formation of a non-productive ESI complex involving Sfβgly, substrate and Tris. In addition, Tris binding reduces by 3 fold the enzyme affinity for the substrate. Hence, at concentrations higher than the *K*_i_, Tris can completely abolish Sfβgly activity, whereas even at lower concentrations the presence of Tris causes underestimation of β-glucosidase kinetic parameters (*K*_m_ and *k*_cat_). In agreement to the inhibition mechanism, computational docking showed that Tris could bind to a pocket placed at the lateral of the active site opening in the Sfβgly-substrate complex, hence leading to the formation of a ESI complex. Computational docking also showed that Tris may find binding spots in the interior of the active site of the Sfβgly and several GH1 β-glucosidases. Moreover, the variety of their active site shapes results in a multiplicity of binding profiles, foreseeing different inhibition mechanism. Thus, Tris inhibition is probably common among GH1 β-glucosidases. This remark should be taken into account in their study, highlighting the importance of the appropriate buffer for accurate enzyme characterization.

## 1. Introduction

Glycoside hydrolases (GH) are enzymes found in all living forms. Based on their sequence and structural similarities, GH enzymes are categorized into 189 families in the CAZY databank (http://www.cazy.org) [1]. The GH1 family includes β-glucosidases (EC 3.2.1.21) which catalyze the hydrolysis of O-or S-glycosidic linkages of β-glycosides, playing important functions in numerous biological processes [2].

The GH1 β-glucosidases present a (β/α)_8_ barrel fold in which the active site is placed among the loops projecting from the C-terminal edge of the β-sheet that forms their core barrel. The active site, shaped like a tunnel or pocket, is schematically divided into subsites, each one corresponding to the set of residues that interact with one monosaccharide unit of the substrate. Hence, at least two subsites are present. The subsite -1 binds the substrate glycone, *i*.*e*. monosaccharide at its non-reducing end, and the subsite +1, where the substrate aglycone is positioned. GH1 β-glucosidases active upon oligocellodextrins usually have several positive subsites (+1, +2, +3 and so on) [3; 4]. Residues forming the subsite -1 are conserved, whereas the positive subsites present a high variability, conferring different shapes and specificities to this active site region [2; 4]. Between the subsites -1 and +1 are placed the residues involved in the hydrolysis reaction, two conserved glutamate residues that act as catalytic acid/base and nucleophile. Additionally, arginine and tyrosine residues may be present modulating the catalytic nucleophile ionization [4; 5].

Tris (2-amino-2-(hydroxymethyl)-1,3-propanediol) is a buffering compound frequently employed in Biochemistry and Molecular Biology [6; 7]. It is a synthetic organic compound with the formula (HOCH_2_)_3_CNH_2_, and is also known as Tris base, THAM, Trometamol and tris(hydroxymethyl)aminomethane. Its inhibitory effect was observed for aminopeptidases, amylases and β-glucosidases [8-14]. However, there have been no efforts made to elucidate the inhibition mechanism through classical enzyme kinetics experiments.

Here we tackled that question by using the recombinant GH1 β-glucosidase from *Spodoptera frugiperda* (Sfβgly). This is a digestive enzyme associated with the glycocalyx of the midgut epithelial cells from the insect *S. frugiperda*, also known as fall armyworm [15]. Sfβgly has already been characterized regarding the preference for the substrate at the subsites -1 and +1, as well as its biophysical properties such as thermal stability and dimerization [15 – 19].

The crystallographic structure of Sfβgly (PDB ID: 5CG0) [17] reveals an active site comprised of two subsites. Subsite −1, which binds the monosaccharide at the non-reducing end of the substrate, is characterized by a collection of residues that form hydrogen-bonds with the ligand [20]. These interactions suggest a putative binding site for small molecules that could also be involved in hydrogen-bonds, like Tris. Indeed, the crystallographic structure of Sfβgly shows a Tris molecule bound into the subsite -1. This finding was unintentional and only occurred because the buffer used in the protein crystallization contained Tris [17]. However, this indicates the potential inhibitory effect of Tris on this enzyme.

Here enzyme kinetic experiments were planned to characterize the mechanism of inhibition of Sfβgly by Tris. Noteworthy, we used Tris concentrations similar to those tipically used in buffers, in the mM tens range. The binding of a Tris molecule to Sfβgly was also tested using computational docking, enabling comparison between these *in silico* models and the crystallographic structure. The kinetic and structural data were combined to produce a coherent view of the interaction between Sfβgly and Tris, as well as the resulting inhibition mechanism. Finally, we discussed the possibility that Tris is a general inhibitor of GH 1 β-glucosidases. This observation should be taken into account in their study, emphasizing the importance of the appropriate buffer for accurate enzyme characterization.

## 2. Material and Methods

### 2.1 Protein preparation

Sfβgly production and purification have been previously reported [17; 18]. The homogeneity of the samples was checked by SDS-PAGE [21]. Protein concentration was measured using the bicinchoninic acid (BCA) assay [22]. The purified Sfβgly sample was submitted to buffer exchange using PD Minitrap G-25 columns (Cytiva, Marlborough, MA, USA). The final sample was then stored in 100 mM phosphate buffer pH 6 at 4°C (Supplementary Figure 1).

### 2.2 Determination of the Sfβgly catalytic activity

The hydrolysis activity of Sfβgly upon *p*-nitrophenyl β-glucoside (NPβglc) and cellobiose (C2) were determined as previously described [23]. Initial rates were calculated from the slope of lines correlating the [product] and time. Linear regression and correlation coefficients (*R*^2^) were used to evaluate the lines. Lines with *R*^2^ value higher than 0.95 were accepted.

Transglycosylation reactions were detected based on the ratio of the two products, *p*-nitrophenolate and glucose, formed from 20 mM NPβglc [23]. The *p*-nitrophenolate and glucose standard curves, prepared with and without Tris, showed no significant differences.

Substrates and Tris were prepared in 100 mM phosphate buffer pH 6. Assays were performed at 30°C.

The effect of Tris on Sfβgly stability was assessed by incubating enzyme samples in the presence and absence of 300 mM Tris for 18 hours at 30°C. Next, Tris was removed by washing the enzyme sample with 50 volumes of 100 mM phosphate buffer pH 6. Centrifugal filter devices (Amicon Ultracel-3K, Millipore, Burlington, MA, USA) were used for this step. Finally, the enzyme activities of those samples after Tris removal were determined using 10 mM NPβglc, as mentioned above.

Finally, to prevent unexpected changes in the pH of the phosphate buffer, Tris was initially combined with buffering components and only after that the pH was set to 6. NaCl was added to the phosphate buffer without Tris to adjust the ionic strength.

### 2.3 Inhibition of the Sfβgly activity with tris

The initial rate of hydrolysis of 10 different substrate concentrations (NPβglc or C2) was determined in the presence of 5 different Tris concentrations (0 – 120 mM). NPβglc ranged from 0.25 to 10 mM, whereas C2 from 0.6 to 16 mM.

Initial rates were determined based on three reactions performed at 30°C. NPβglc, C2 and Tris were prepared in 100 mM phosphate buffer pH 6.0. Products, *p*-nitrophenolate and glucose, were detected as previously described [23]. NaCl was added to adjust the ionic strengths among reactions, with 120 mM Tris as the reference. Experiments were performed with two different enzyme samples. Data were analyzed using Lineweaver–Burk plots. Linear fittings were accepted when showing *R*^2^ higher than 0.9. *K*_i_ and α determination were based on procedures appropriated to the inhibition mechanism [24] and expressed as median and standard deviation.

### 2.4 Computational docking between Tris and GH1 β-glucosidases

GH1 β-glucosidases spatial coordinates were obtained from the PDB Data Bank. Entries were indicated in the text and figure legends. Tris coordinates were generated using Gabedit v.2.5.1 [25]. AutoDockTools 1.5.7 was used to delete water molecules, ions and other molecules and add hydrogen atoms in the protein structures, and build the grid box coordinates covering all protein atoms [26]. Dockings were performed using AutoDock Vina [27] searching the 9 best models. Tris was docked in the mono-protonated state (+1 charge). The Sfβgly-NPβglc complex coordinates were obtained by structural alignment between the structures 5CG0 and 3AI0. After the alignment a PDB file was built by retaining only the coordinates of the Sfβgly (5CG0) and the NPβglc (from the 3AI0 file). The Tris-enzyme complexes representing the different docking solutions were visualized by using PYMOL v0.99 software (Schrodinger LLC, NewYork, NY, USA).

## 3. Results and Discussion

The addition of 40 mM Tris reduced to 60% the initial rate of NPβglc hydrolysis catalyzed by Sfβgly (Fig. 1). However, Tris did not irreversibly inactivate Sfβgly, as pre-incubation with 300 mM Tris for 18 h did not change the reaction rate in the assays performed after Tris removal (Fig. 1). Furthermore, Tris did not favor the occurrence of transglycosylation reaction catalyzed by Sfβgly given that in the presence of 30 mM Tris the ratio of glucose to *p*-nitrophenolate formation from NPβglc is 1 (Fig. 1). As known, in transglycosylation reactions having NPβglc as substrate, the glucose is incorporated in the product, so that ratio should drop. Importantly standard curves for *p*-nitrophenolate and glucose prepared in the presence and absence of 120 mM Tris demonstrated that it does not affect the detection of these products (Fig. 1). In short, these results suggest that Tris act as a reversible inhibitor of Sfβgly.

**Figure 1.**
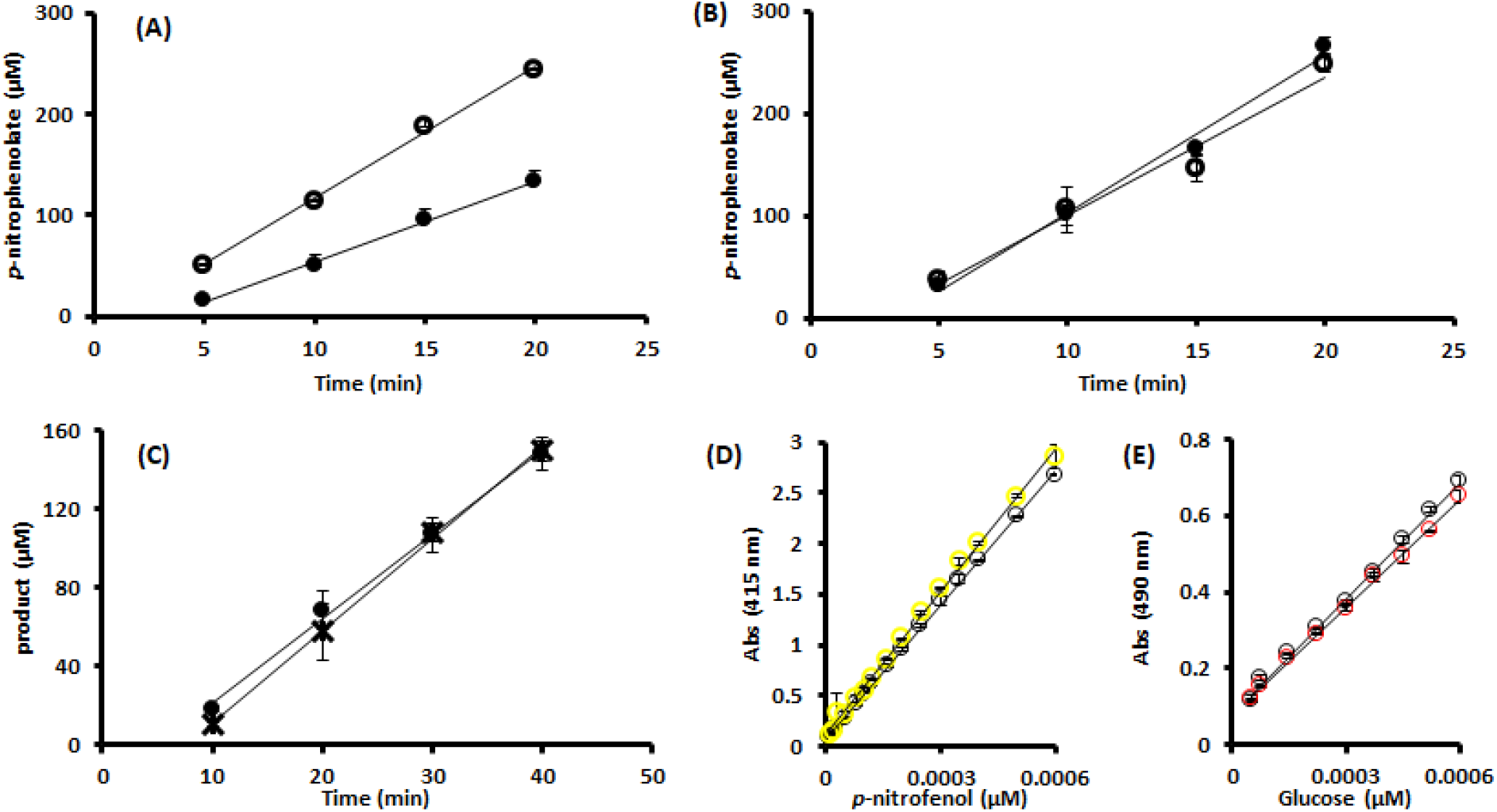
Effect of Tris on the Sf⍰gly activity and stability. (**A**) Initial rate of 10 mM NP⍰glc hydrolysis catalyzed by Sfβgly in the absence (○; 13 μM.min^−1^) and presence of 40 mM Tris (•; 7.9 μM.min^−1^). (**B**) Initial rate of 10 mM NP⍰glc hydrolysis catalyzed by Sfβgly previously incubated with 300 mM Tris at 30 °C for 18 h (•; 15 μM.min^−1^) and without Tris (○; 13 μMmin^−1^). Both experiments, shown in A and B, were performed with 0.09 μM Sfβgly. Data are mean and standard deviation of three determinations of the product formed in each incubation time using three separate assays with the same enzyme sample. The substrate was prepared in 100 mM phosphate buffer pH 6.0. Activity assays were performed at 30 °C. (**C**) Tests aiming at detection of the transglycosylation reaction catalyzed by Sfβgly (0.009 μM). Production of *p*-nitrophenolate (○) and glucose (**\***) from 20 mM NP⍰glc in the presence of 30 mM Tris. The substrate was prepared in 100 mM phosphate buffer pH 6.0. Activity assays were done at 30 °C. Data are mean and standard deviation of three determinations of the product formed in each incubation time using the same enzyme sample. (**D**) Standard curve of *p*-nitrophenolate with (○) and without (**○**) 120 mM Tris. Slopes are 4420 and 4721 Abs_415nm_. μM^-1^, respectively. (**E**) Standard curve of glucose with (○) and without (**○**) 120 mM Tris. Slopes are 1007 and 935 Abs_415nm_. μM^-1^, respectively. *R*^2^ are higher than 0.99.

To uncover the mechanism of that inhibition, the initial hydrolysis rate of various NPβglc concentrations determined at different Tris concentrations were analyzed using Lineweaver-Burk plots. The experiment was performed three times with two independent enzyme samples (Supplementary Figure 1), producing the same pattern (Fig. 2; Supplementary Figures 2 and 3). Increments of Tris concentration generated a set of lines with increasing *K*_s_/*k*_3_ (slope) and 1/*k*_3_ (intercept). In addition, those lines intersected above the 1/[S] axis in the second quadrant. Finally, the apparent *K*_s_/*k*_3_ and apparent 1/*k*_3_ showed a linear relation with the Tris concentration (Fig. 2; Supplementary Figures 2 and 3). These features define a linear mixed-type inhibition mechanism, specifically an intersecting, linear, noncompetitive inhibition (Fig. 2) [24].

**Figure 2.**
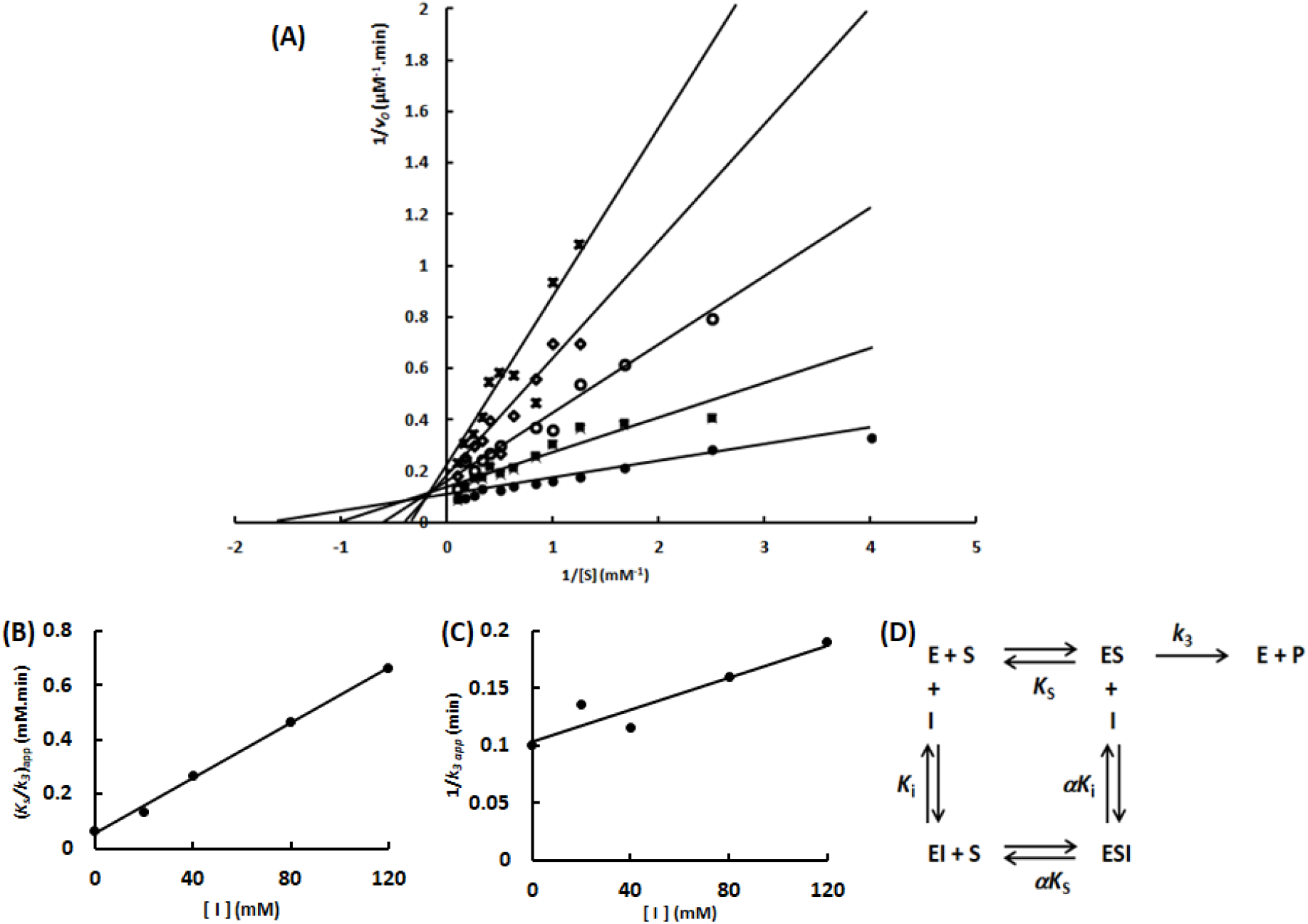
Characterization of the Tris inhibition mechanism upon Sfβgly. (**A**) Lineweaver-Burk plots showing the effect of different Tris concentrations (•, 0; ◼, 20 mM; ○, 40 mM; ◊, 80 mM; **x**, 120 mM) on the initial rate of hydrolysis of the substrate NPβglc. (**B**) Tris effect on the apparent *K*_s_/*k*_3_ (calculated from the line slope). (**C**) Tris effect on the apparent 1/ *k*_3_ (calculated from the line intercept). NPbglc and Tris were prepared in 100 mM phosphate buffer pH 6.0. Rates were determined at 30°C. Rates are mean of three product determination using the same enzyme sample. Experiment performed with the enzyme sample #1. Independent experiments were performed using two different enzyme sample (Supplementary Figures 1 and 2). (**D**) Linear mixed type inhibition mechanism (intersecting, linear, noncompetitive). S, substrate NPβglc; I, inhibitor Tris; E, enzyme Sfβgly; P, product; *K*_s_, dissociation constant for the ES complex; *K*_i_, dissociation constant for the EI complex; *k*_3_, rate constant for product formation; α, factor that represents the mutual hindering effect between S and I (α >1) [24].

A linear mixed-type inhibition mechanism indicates that Tris (inhibitor) and NPβglc (substrate) may both bind to the Sfβgly enzyme, forming a non-productive complex ESI. So even in the presence of a theoretical infinite substrate concentration, which should result in the limiting initial rate (*V*_max_), the presence of Tris would drive a fraction of the enzyme population to be locked in the inactive ESI complex. Hence, a lower apparent *V*_max_ is reached in the presence of Tris. As a result, the line intercept, *i*.*e*. 1/*k*_3_ which is proportional to 1/*V*_max_, (Fig. 2; Supplementary Figures 2 and 3) increases with increasing Tris concentration.

Besides that, both the inhibitor Tris and the substrate NPβglc, can bind to the free enzyme and the ES complex (Fig. 2). However, the presence of one ligand, substrate or inhibitor, hampers the binding of the second on. Hence, the affinity between substrate and EI complex is lower than that between substrate and E, whereas the inhibitor affinity for the ES complex is also lower than for E. Therefore, α*K*_i_ and α*K*_s_ are higher than *K*_i_ and *K*_s_, respectively. Consequently, the factor α, which is higher than 1, represents the detrimental effect that the first ligand (substrate or Tris) exerts upon the second ligand binding.

The same experiment was repeated using C2, a natural substrate of Sfβgly, and two independent enzyme samples. The same inhibitory mechanism was observed (Fig. 3; Supplementary Figure 4).

Then, considering that Tris was shown to be a linear mixed-type inhibitor of Sfβgly, the apparent *K*_s_/*k*_3_ *versus* [Tris] and apparent 1/*k*_3_ *versus* [Tris] (Fig. 2; Fig. 3; Supplementary Figures 2 - 4) plots were employed to calculate the *K*_i_ and α (Table 1) [24].

**Table 1.**
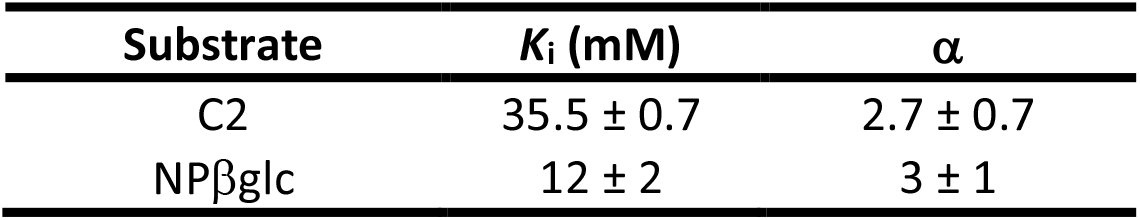
Parameters of the Tris inhibition mechanism upon the β-glucosidase Sfβgly.

**Figure 3.**
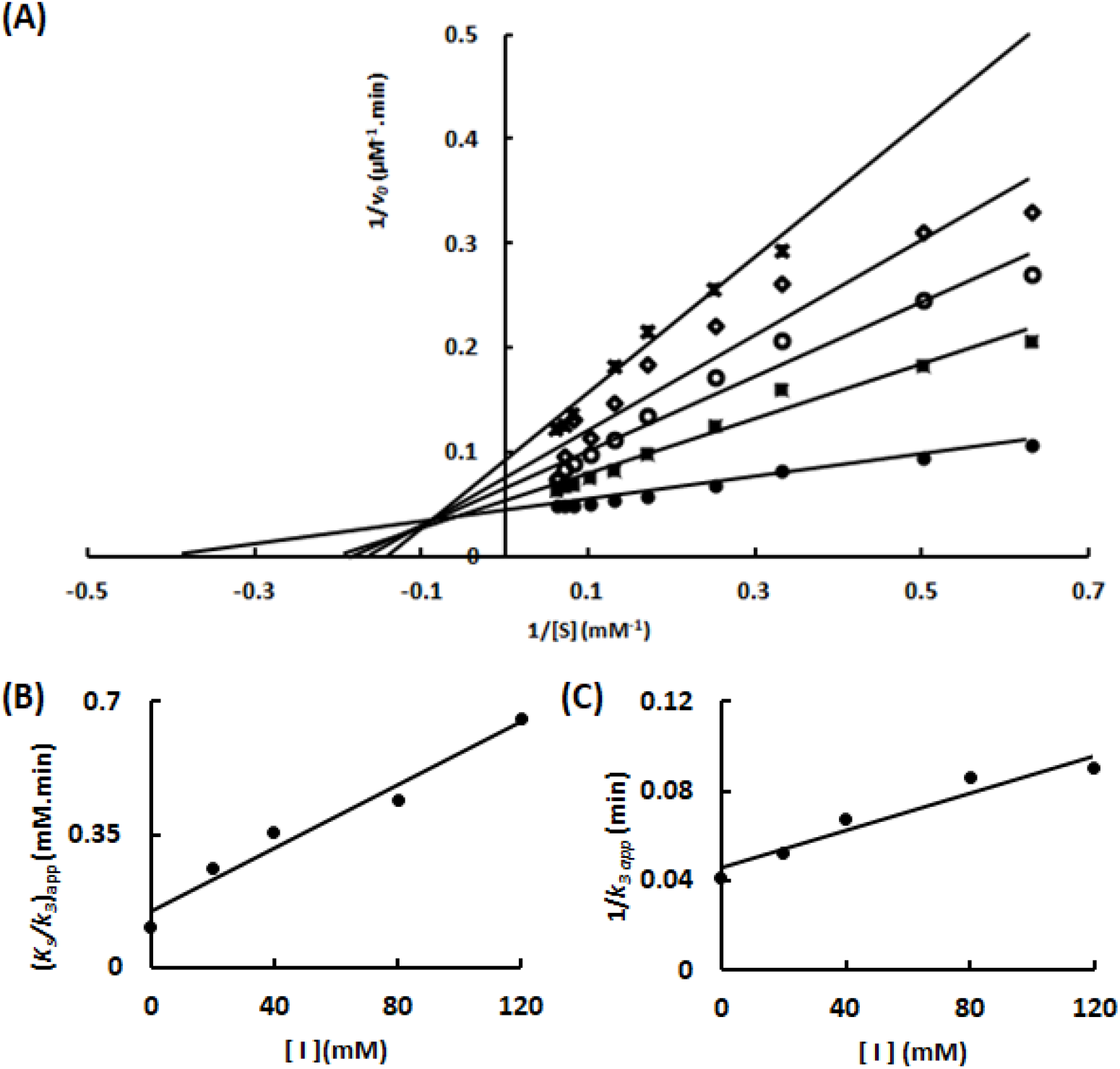
Characterization of the Tris inhibition mechanism upon hydrolysis of the substrate C2 catalyzed by Sfβgly. (**A**) Lineweaver-Burk plots showing the effect of different Tris concentrations •, 0; ◼, 20 mM; ○, 40 mM; ◊, 80 mM; **x**, 120 mM) on the initial rate of C2 hydrolysis. (**B**) Tris effect on the apparent *K*_s_/*k*_3_ (calculated from the line slope). (**C**) Tris effect on the apparent 1/ *k*_3_ (calculated from the line intercept). C2 and Tris were prepared in 100 mM phosphate buffer pH 6.0. Rates were determined at 30°C. Rates are mean of three product determination using the same enzyme sample. Two independent experiments were performed using two different enzyme samples (Supplementary Figure 4).

The inhibition mechanism is shown on Fig. 2. *K*_i_ represents the dissociation constant of the Sfβgly enzyme-Tris complex (EI). α corresponds to the mutual hindering effect involving substrate and Tris. Data are averages and standard deviation. n = 2 for C2 and n = 3 for NPβglc.

The similar *K*_i_ observed with two different substrates suggests that Tris is binding in the same site in both cases. The factor α > 1 confirms a significant mutual impediment involving Tris and the substrate, showing that the first ligand causes a 3-fold decrease in the affinity of the second one. The physical nature of this effect is not clear by now, but a steric hindrance resulting in alteration of the distances and angles of the non-covalent interactions involved in the substrate or inhibitor binding could be proposed.

Finally, the previous suggestion of non-specific binding of Tris to Sfβgly implying no inhibitory effect [28] was not supported by the results presented here.

The computational docking of a Tris molecule into the Sfβgly crystallographic structure (5CG0; [17]) revealed a set of similarly possible binding sites (Supplementary Table 1; Supplementary Figure 5) within the active site pocket encircled (3.5 Å) by the subsite -1 residues Y331, W444 and E451 and around the catalytic residues E187 and E399 (Fig. 4). Tris had been previously observed in the Sfβgly crystallographic structure, also within that same site (Fig. 4) [5; 15; 16; 17; 20].

**Figure 4.**
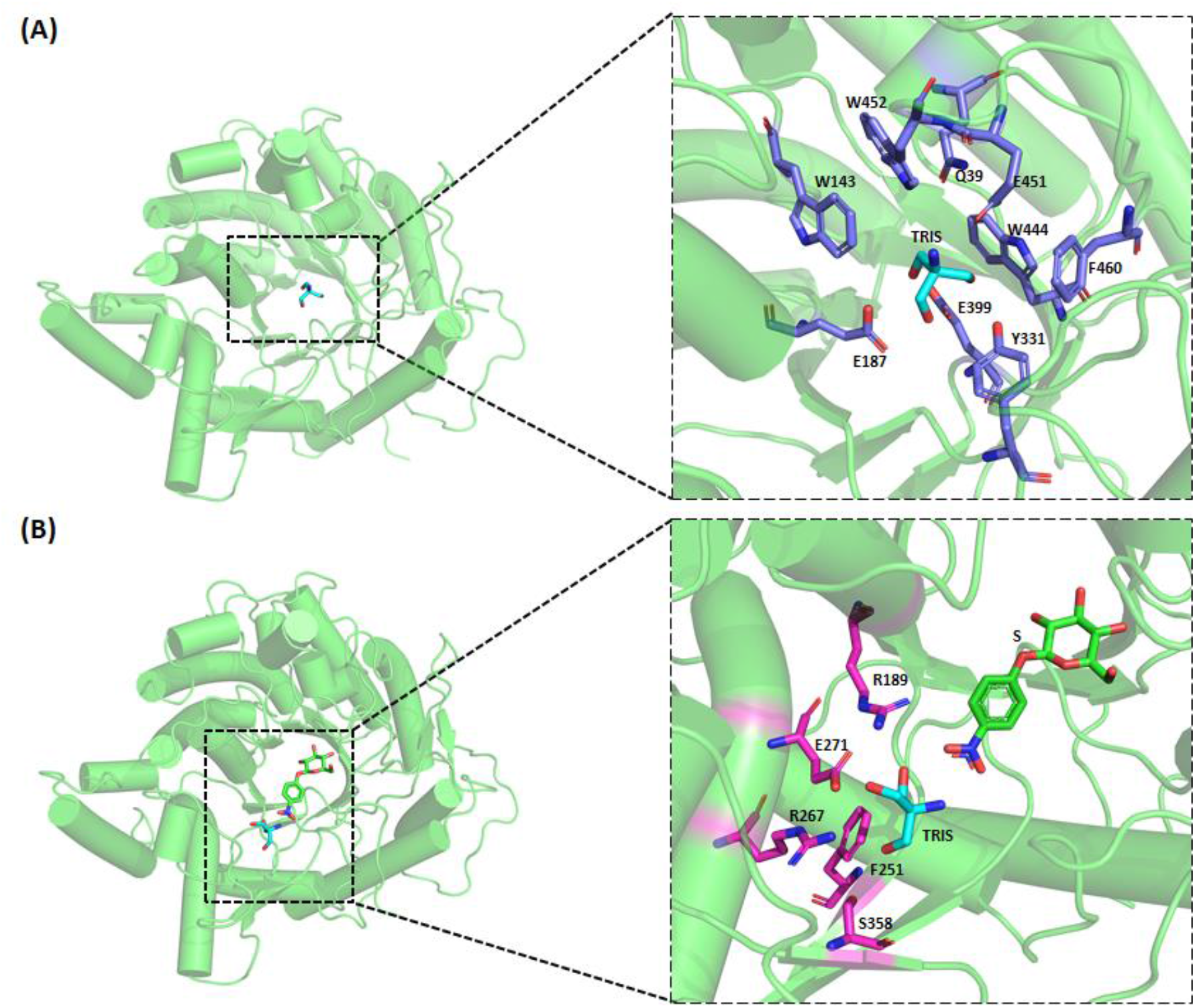
Tris binding in the β-glucosidase Sfβgly and the Sfβgly-NPβglc complex. (**A**) Binding spots in the free enzyme revealed by computational docking (solutions #1 to #7 – Supplementary Table 1; Supplementary Figure 5). Residues interacting with Tris are shown in blue. (**B**) Binding spots in the enzyme-substrate complex revealed by computational docking (Supplementary Table 2; Supplementary Figure 6) Residues showed in the detailed structures (panels A and B) are within 3.5 Å of the Tris molecule.

But the above binding mode is not compatible with the mixed inhibitory mechanism observed here (Fig. 2). Actually, it would be consistent to a simple competitive inhibition [24], as suggested before [14].

On the other side, when the Sfβgly-NPβglc complex (ES) was employed in the molecular docking (Supplementary Table 2; Supplementary Figure 6), a solution showed the Tris molecule placed in a lateral pocket in the active site opening, surrounded by (3.5 Å) residues R189, F251, R267, E271 and S358 (Fig. 4). Imidazole had been previously found to inhibit Sfβgly by interacting in that same region [23]. Hence, Tris positioning in such pocket could be compatible with the simultaneous binding of the substrate in the -1 and +1 subsites. However, their simultaneous presence could misplace the substrate relatively to the catalytic residues, which would produce a non-productive ESI complex. Therefore, such Tris docking solution (Fig. 4) could generate the linear mixed inhibition mechanism observed here (Fig. 2).

The observation of two putative binding sites for Tris in the Sfβgly active site is not incoherent since Tris has multiple groups that could be involved in hydrogen bonds (3 hydroxyls and 1 amine) with the many polar residues along the active site tunnel, which usually interact with the hydroxyl groups of the Sfβgly natural substrates, *i*.*e*. oligocellodextrins [5; 16; 20].

Indeed, based on this, it could be anticipated that small molecules presenting hydrogen bond potential (for instance Tris, imidazole, glycerol and so on) may find binding spots along the active site of the GH1 β-glucosidases. Computational docking of Tris into several GH1 β-glucosidases supports that hypothesis (Supplementary Figure 7). Interestingly, the shape variability of their active site, particularly in the more external portions that probably constitutes the positive subsites and opening, suggest a multiplicity of possible binding modes, which could produce unlike inhibition mechanisms for different β-glucosidases. Thus, Tris inhibition is probably common among GH1 β-glucosidase. This observation should be considered in the study of GH 1 β-glucosidases, stressing the importance of the appropriate buffer for accurate enzyme characterization.

## Abbreviations

Sfβgly: GH1 β-glucosidase from *Spodoptera frugiperda* (PDB 5CG0);
NPβglc: *p*-nitrophenyl β-glucoside;
C2: cellobiose;
Tris: 2-amino-2-(hydroxymethyl)-1,3-propanediol

## Author Contributions

**Rafael S. Chagas:** conceptualization; formal analysis; investigation; writing original draft; writing review and editing. **Sandro R. Marana:** conceptualization; funding acquisition; resources; formal analysis; investigation; writing - original draft; writing - review and editing.

## Supplementary Materials

**Supplementary Figure 1.**
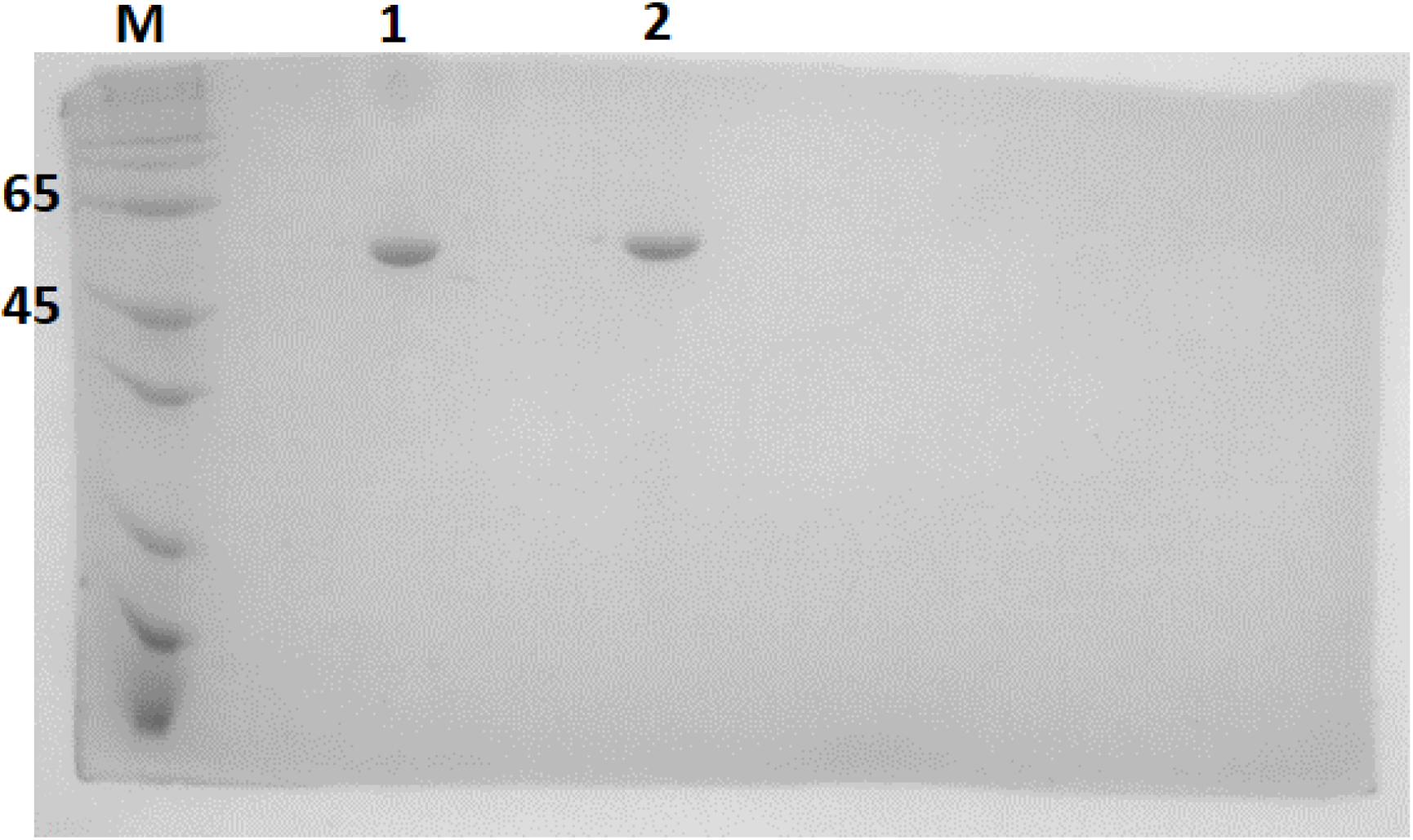
SDS-PAGE of the purified recombinant Sfβgly. Lanes: M – Molecular weight marker (kDa); 1 – Sfβgly sample 1; 2 - Sfβgly Sample 2.

**Suplementary Figure 2.**
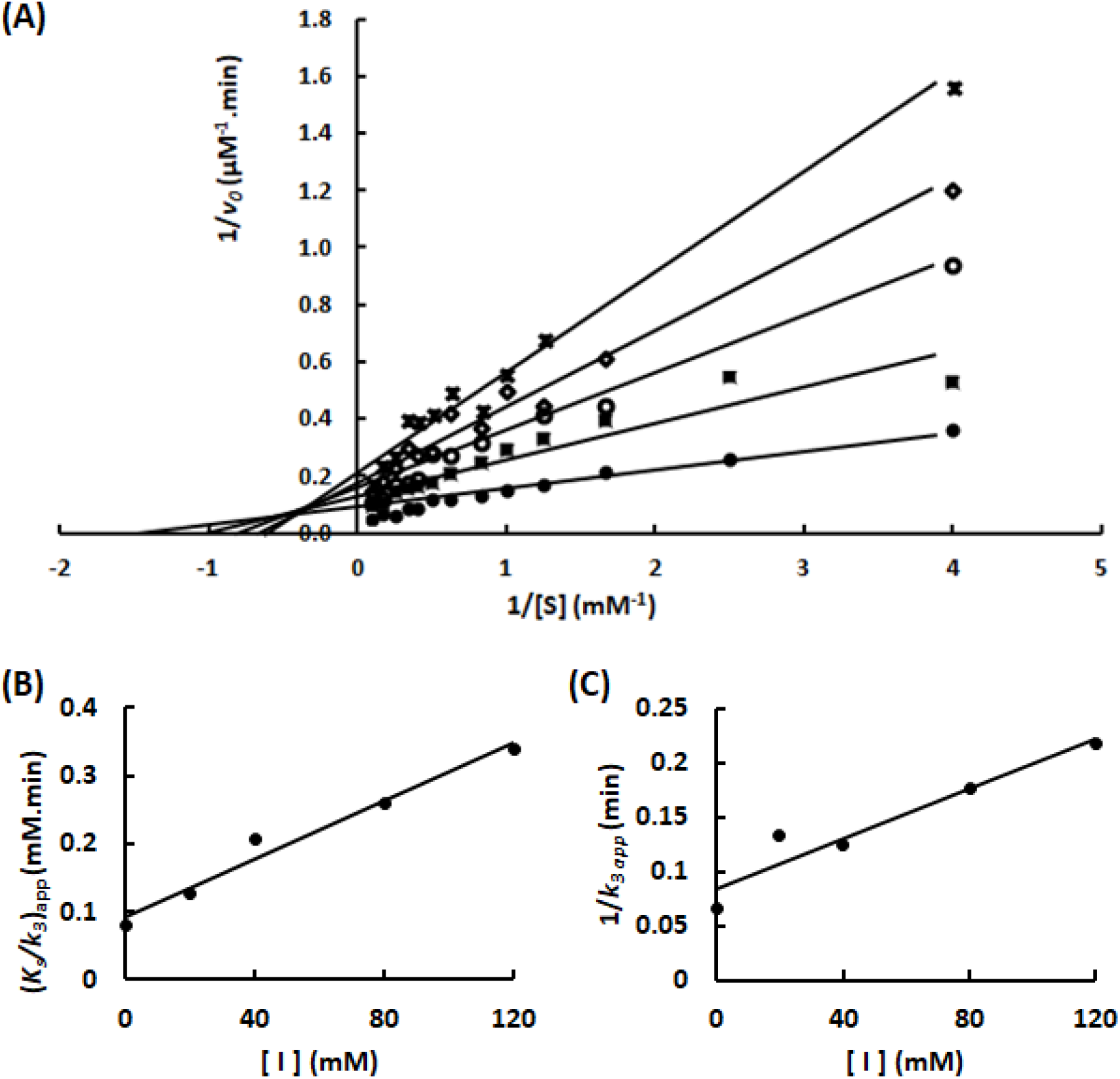
Characterization of the Tris inhibition mechanism upon Sfβgly. Enzyme sample #1. Substrate NPβglc. Characterization of the Tris inhibition mechanism upon Sfβgly. (**A**) Lineweaver-Burk plots showing the effect of different Tris concentrations (•, 0; ◼, 20 mM; ○, 40 mM; ◊, 80 mM; **x**, 120 mM) on the initial rate of hydrolysis of the substrate NPβglc. (**B**) Tris effect on the apparent *K*_s_/*k*_3_ (calculated from the line slope). (**C**) Tris effect on the apparent 1/ *k*_3_ (calculated from the line intercept). NPβglc and Tris were prepared in 100 mM phosphate buffer pH 6.0. Rates were determined at 30°C. Rates are mean of three product determination using the same enzyme sample. Duplicate, enzyme sample #1.

**Supplementary Figure 3.**
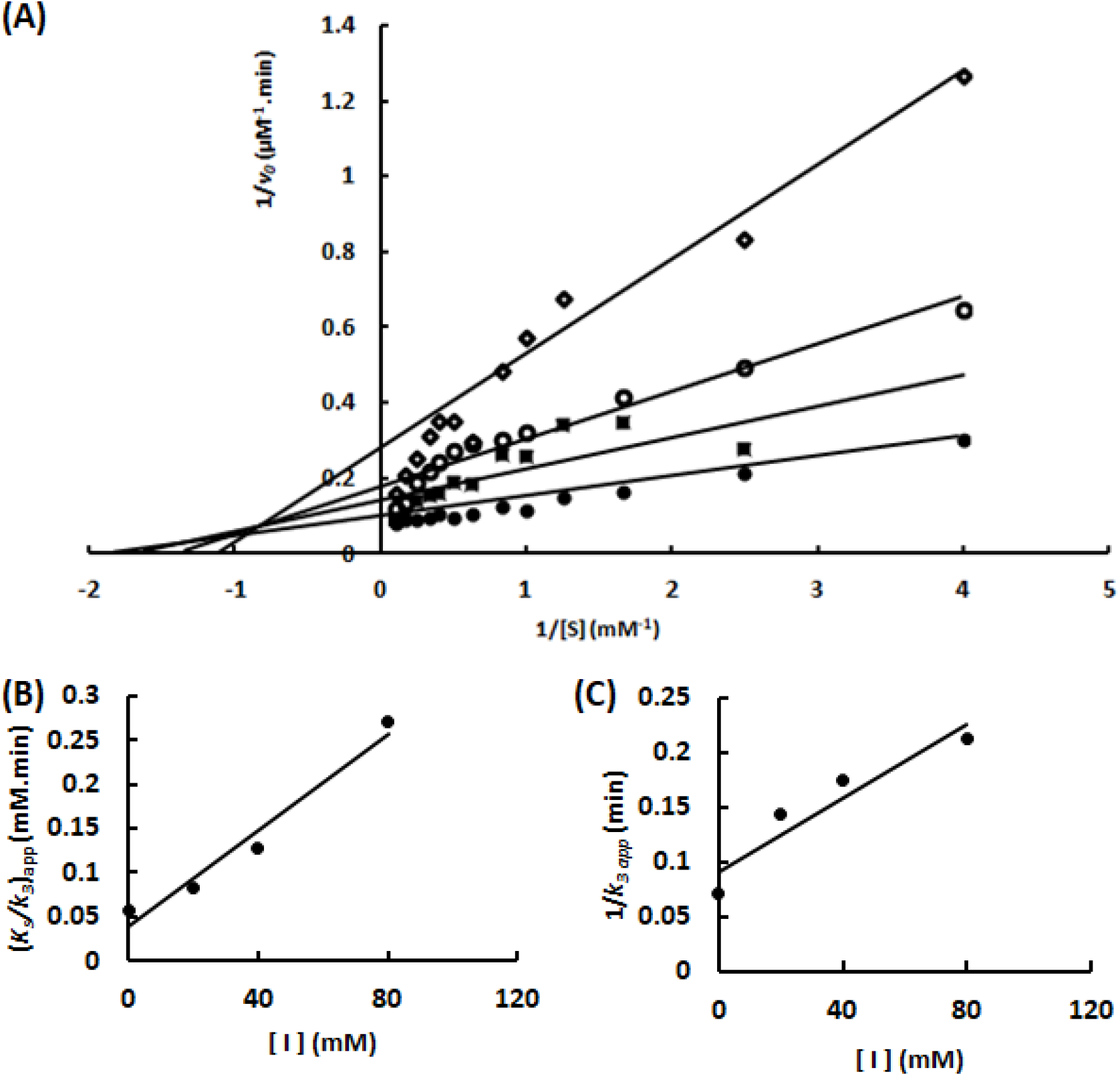
Characterization of the Tris inhibition mechanism upon Sfβgly. Enzyme sample #2. Substrate NPβglc. Characterization of the Tris inhibition mechanism upon Sfβgly. (**A**) Lineweaver-Burk plots showing the effect of different Tris concentrations (•, 0; ◼, 20 mM; ○, 40 mM; ◊, 80 mM; **x**, 120 mM) on the initial rate of hydrolysis of the substrate NPβglc. (**B**) Tris effect on the apparent *K*_s_/*k*_3_ (calculated from the line slope). (**C**) Tris effect on the apparent 1/ *k*_3_ (calculated from the line intercept). NPβglc and Tris were prepared in 100 mM phosphate buffer pH 6.0. Rates were determined at 30°C. Rates are mean of three product determination using the same enzyme sample. Enzyme sample #2.

**Supplementary Figure 4.**
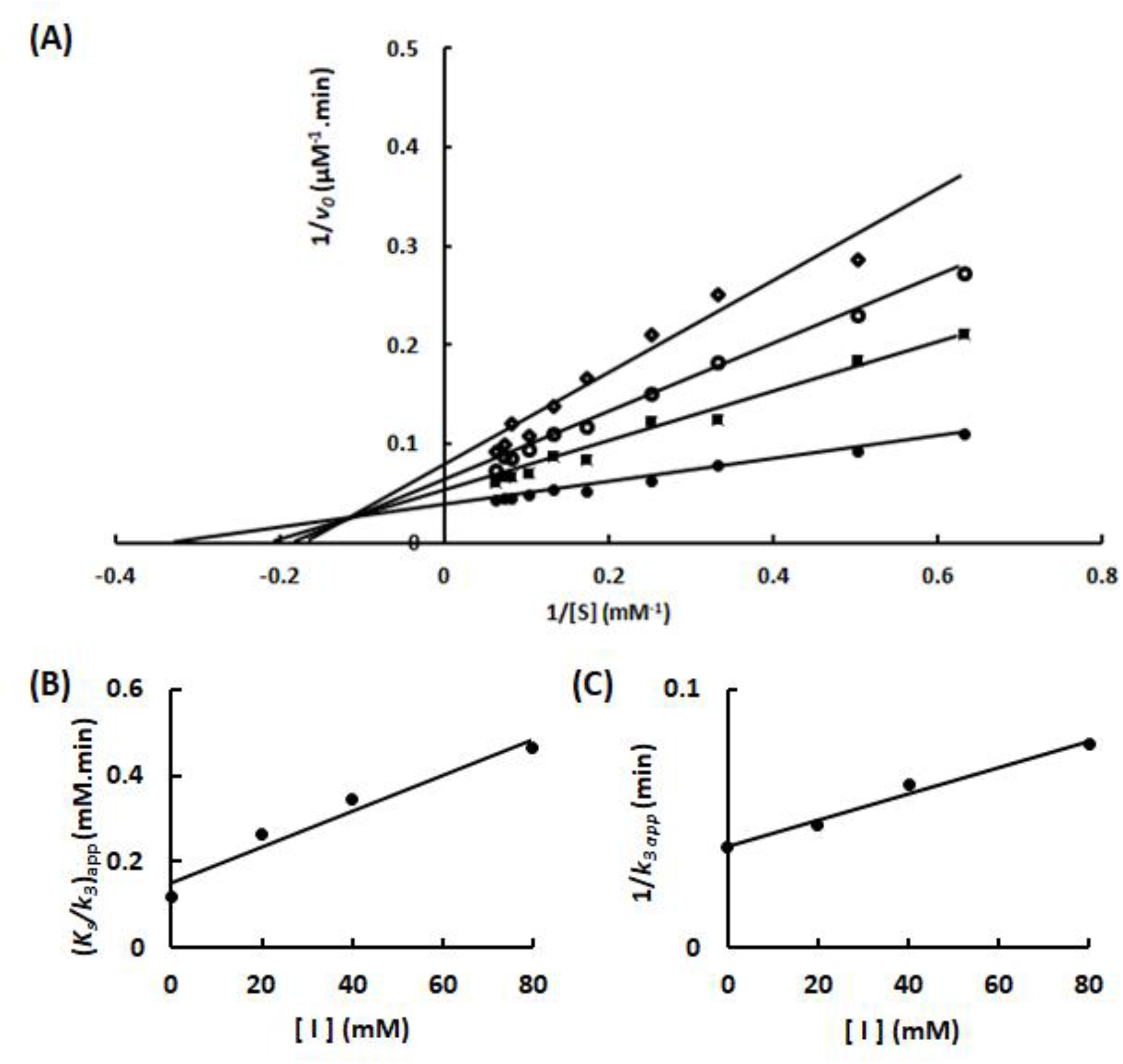
Characterization of the Tris inhibition mechanism upon Sfβgly. Enzyme sample #2. Substrate C2. **Figure 3** – Characterization of the Tris inhibition mechanism upon hydrolysis of the substrate C2 catalyzed by Sfβgly. (**A**) Lineweaver-Burk plots showing the effect of different Tris concentrations •, 0; ◼, 20 mM; ○, 40 mM; ◊, 80 mM; **x**, 120 mM) on the initial rate of C2 hydrolysis. (**B**) Tris effect on the apparent *K*_s_/*k*_3_ (calculated from the line slope). (**C**) Tris effect on the apparent 1/ *k*_3_ (calculated from the line intercept). C2 and Tris were prepared in 100 mM phosphate buffer pH 6.0. Rates were determined at 30°C. Rates are mean of three product determination using the same enzyme sample. Enzyme sample #2.

**Supplementary Table 1.** Docking solutions for Tris binding in the GH1 β-glucosidase Sfβgly.

| Autodock solution | Binding score | Sfβgly residue within 3.5 Å of the Tris molecule | Sfβgly region |
| --- | --- | --- | --- |
| 1 | -4.6 | E187, Y331, E399, W444, E451 | Active site; -1 subsite |
| 2 | -4.5 | Q39, E187, Y331, E399, W444, E451, W452 | Active site; -1 subsite |
| 3 | -4.5 | W143, E187, E399, W444, E451, W452, F460 | Active site; -1 subsite |
| 4 | -4.2 | W143, E187, Y331, E399, W444, E451 | Active site; -1 subsite |
| 5 | -4.1 | E187, E399, W444, E451 | Active site; -1 subsite |
| 6 | -3.9 | E187, Y331, W444, E451, W452 | Active site; -1 subsite |
| 7 | -3.8 | E187, Y331, E399, W444 | Active site; -1 subsite |
| 8 | -3.7 | N70, D72, I73, A74, Y456, R472 | Surface |
| 9 | -3.6 | N70, D72, I73, A74, Y456 | Surface |

**Supplementary Table 2.** Docking solutions for Tris binding in the Sfβgly-NPβglc complex.

| Autodock solution | Binding score | Sfβgly residue within 3.5 Å of the Tris molecule | Sfβgly region |
| --- | --- | --- | --- |
| 1 | -4.1 | N70, D72, I73, A74, Y456, F467, R472 | Surface |
| 2 | -3.9 | N70, D72, I73 | Surface |
| 3 | -3.8 | N70, D72, I73, A74, R472 | Surface |
| 4 | -3.8 | N70, D72, I73 | Surface |
| 5 | -3.8 | R189, F251, R267, E271, S358 | Pocket |
| 6 | -3.8 | N70, I75, A74, Y456, F467, R472 | Surface |
| 7 | -3.8 | E406, S411, R459, E464, V465 | Surface |
| 8 | -3.7 | N70, D72, I73, A74, F467, R472 | Surface |
| 9 | -3.7 | E406, R459, V465, D466, F467 | Surface |

**Supplementary Figure 5.**
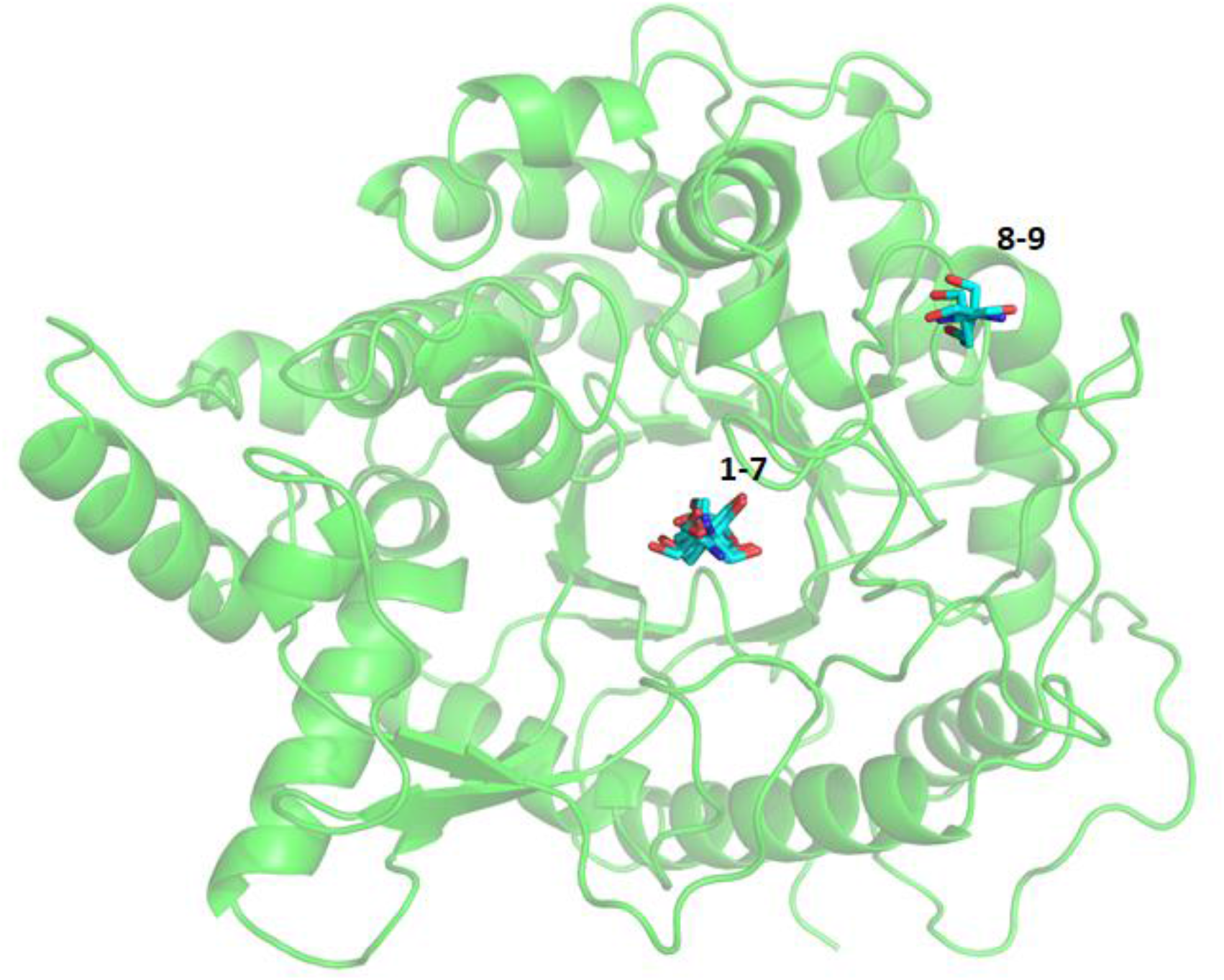
Mapping of the docking solutions for Tris binding in the GH1 β-glucosidase Sfβgly.

**Supplementary Figure 6.**
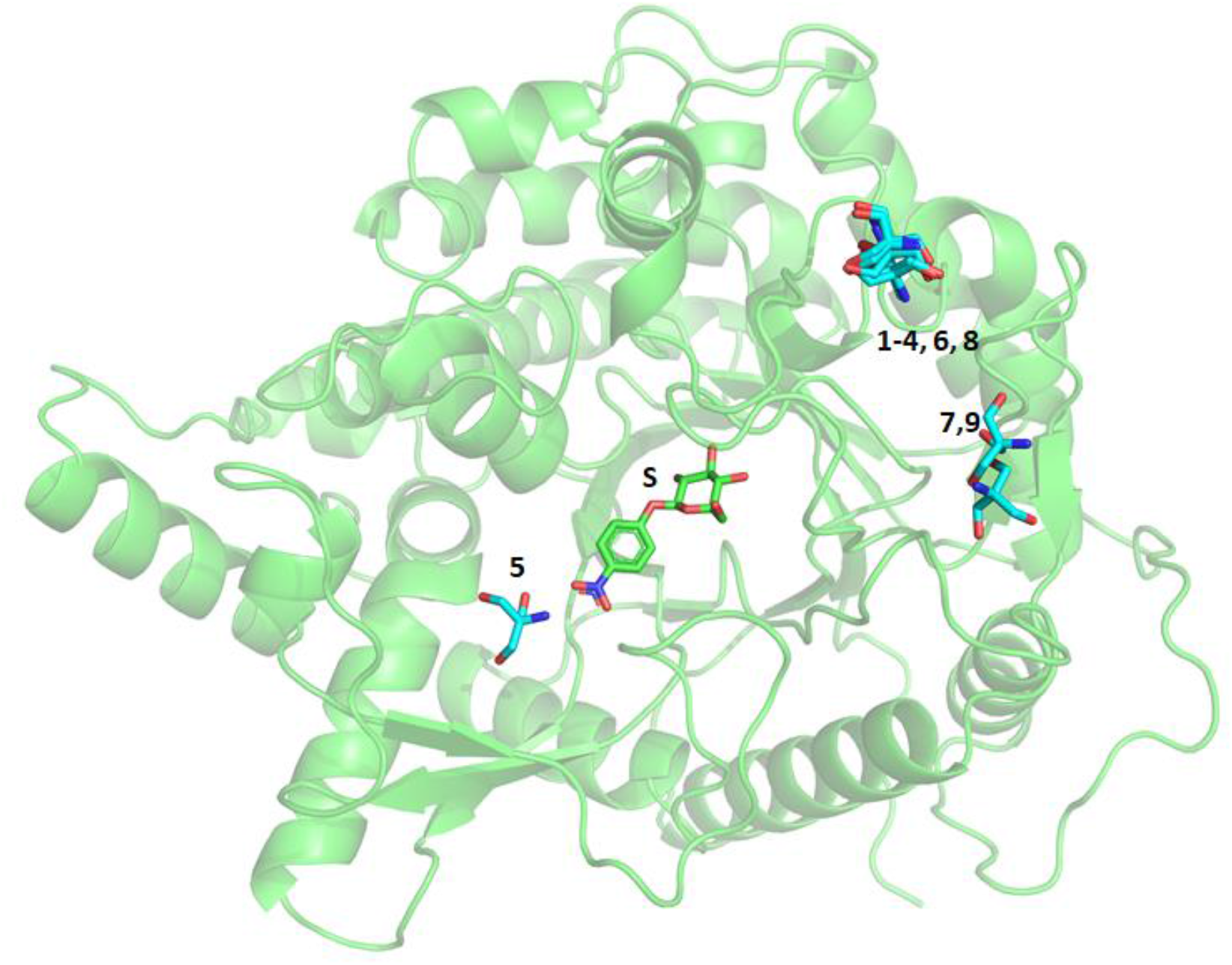
Mapping of the docking solutions for Tris binding in the Sfβgly-NPβglc complex.

**Supplementary Figure 7.**
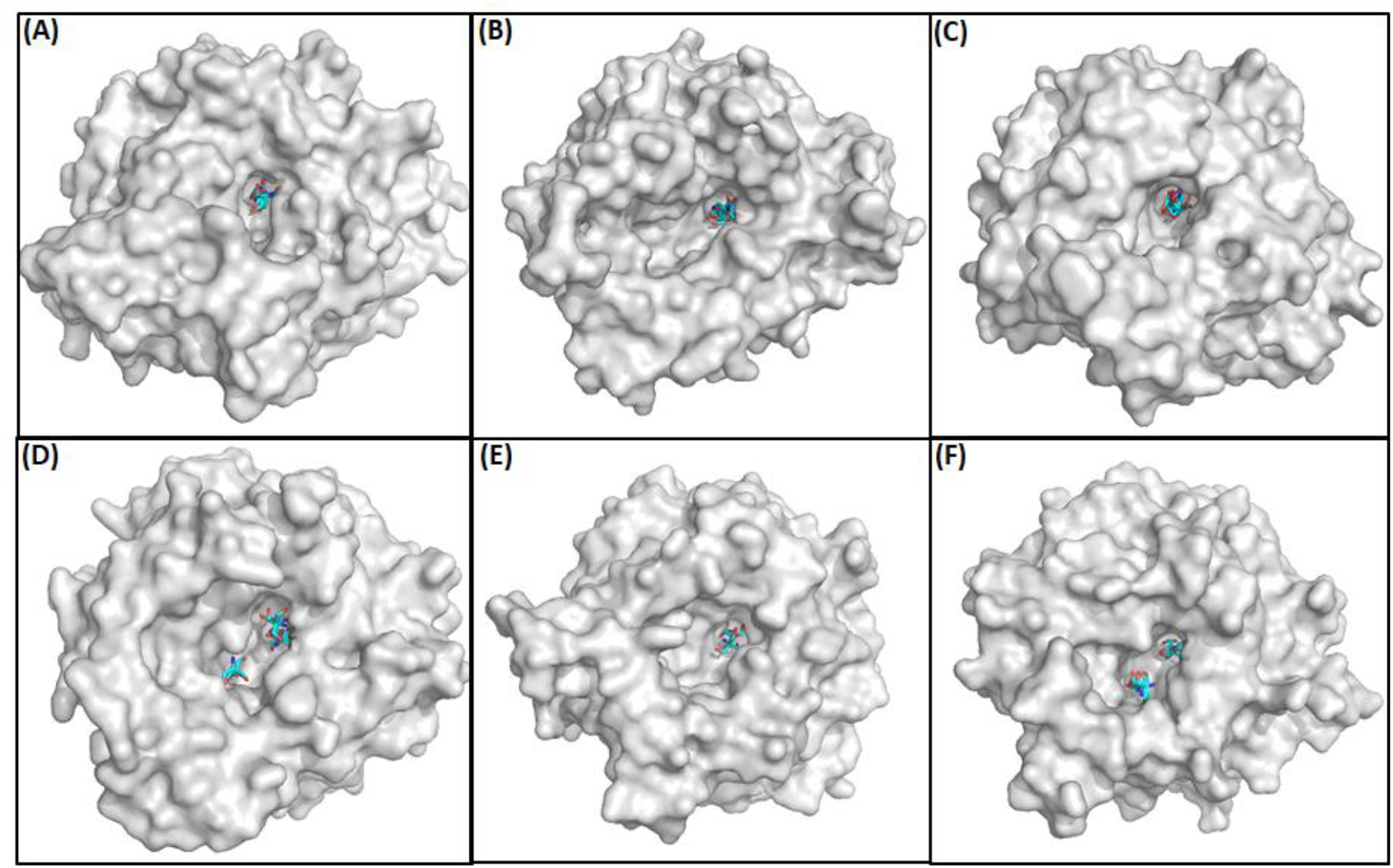
Docking solutions for Tris binding in the active site of different GH1 β-glucosidases. Docking solutions for Tris binding in the active site of different GH1 β-glucosidases. (**A**) 5CG0, Sfbgly; (B) 3GNO from *Oryza sativa*, (C) 1VFF from *Pyrococcus horikoshii*, (D) 5JBO from *Trichoderma harzianum*, (E) 3AHZ from *Neotermes koshunensis*, (F) 2O9P from *Paenibacillus polymyxa*.

